# React-to-Me: A Conversational Interface for Interactive Exploration of the Reactome Pathway Knowledgebase

**DOI:** 10.64898/2025.12.12.693752

**Authors:** Helia Mohammadi, Fatemeh Almodaresi, Gregory F J Hogue, Adam Wright, Marija Orlic-Milacic, Nancy T Li, Amin Mawani, Lincoln Stein

## Abstract

The Reactome Pathway Knowledgebase (www.reactome.org) provides expert-curated information on human biological pathways, molecular interactions, and disease mechanisms. However, its complex data model and keyword-based search interface present accessibility barriers for non-expert users. In contrast, general-purpose conversational AI systems offer intuitive natural language interfaces but lack the domain-specificity, transparent sourcing, and factual reliability required for scientific applications.

To address this gap, we developed React-to-Me (https://reactome.org/chat), a domain-specific conversational assistant that enables users to query Reactome using natural language while maintaining scientific rigor and source traceability. React-to-Me integrates hybrid retrieval-augmented generation (RAG) with constrained language model generation to ensure that all responses are grounded in curated Reactome content and directly linked to corresponding knowledgebase entries. When internal coverage is insufficient, the system defers to trusted external biomedical sources rather than generating speculative or unverified content.

Computational benchmarking confirmed that combining semantic vector search with keyword-based matching substantially improved contextual grounding and factual precision relative to dense-only retrieval baselines. In blinded expert evaluations, grounded responses were more likely to receive higher quality ratings than ungrounded counterparts, with significant gains in factual accuracy, biological specificity, and mechanistic depth. User surveys further indicated strong satisfaction with ease of use, citation reliability, and factual accuracy.

These findings demonstrate that domain-specific grounding can markedly improve the reliability and usability of conversational AI for biological knowledge exploration. React-to-Me provides a transparent and scientifically robust interface for accessing and exploring Reactome content and is freely available at https://reactome.org/chat.

**Graphical Abstract:** 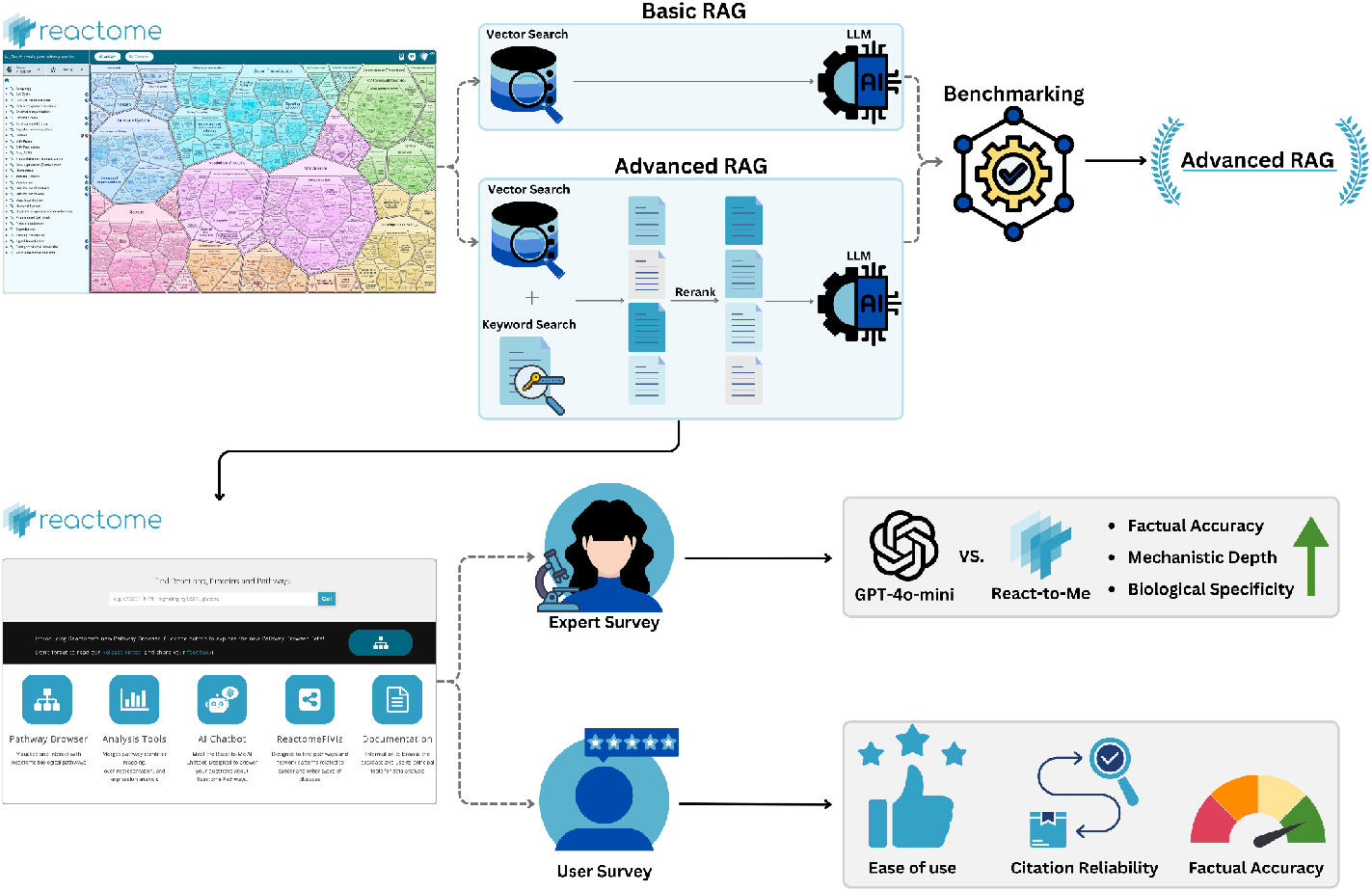

## Introduction

Curated biological knowledgebases are essential resources for the biomedical research community [1–3]. Powered by teams of expert curators, these resources extract, interpret and organize the vast corpus of scientific literature into coherent, structured knowledge. This curated content provides a reliable foundation for downstream analyses that enable data interpretation, guide hypothesis generation, and facilitate the discovery of mechanistic insights. A prime example of a curated biological knowledgebase is Reactome, currently the world’s largest open-access repository of human biological pathways and reactions [4–7]. Reactome is a manually curated and peer-reviewed knowledgebase that captures a detailed hierarchy of molecular entities, their roles in biochemical reactions, and their associated biological pathways. It provides both a browser interface for human users and a RESTful API that supports online and third-party analytic tools such as gene set enrichment and pathway analysis. These capabilities, combined with its open-source and open-access policies, have made Reactome a foundational tool for the molecular biology and disease research communities, serving requests from over 250,000 unique IP addresses per month [6–9].

However, as Reactome has grown in scope and complexity, navigating its content has become increasingly challenging. The current search interface relies on a legacy keyword-based retrieval system that depends on exact term matching rather than conceptual understanding. As a result, queries that omit specific nomenclature, or use alternate phrasing often fail to retrieve relevant information while semantically broad terms can return large sets of isolated and tangentially related entries. For example, a query such as “What genes increase the risk for cancer?” may provide numerous entity-level hits but offers no synthesized explanation of how these genes interact within pathways or contribute to disease processes. Effective use of this interface therefore requires familiarity with Reactome’s domain-specific terminology and its underlying data model [3,10]. This presents a challenge for non-specialists, early-career researchers, clinicians, educators and lay users. To better serve its diverse user community, Reactome needs an interface capable of interpreting user intent, resolving biological terminology, and assembling its curated data into coherent, context-aware summaries.

Recent advances in artificial intelligence, particularly large language models (LLMs) and retrieval-augmented generation (RAG), offer a promising avenue for improving the accessibility of biological knowledgebases like Reactome [11–19]. Unlike traditional search engines, LLMs support dialogue-driven, context-aware exploration, allowing users to interact with complex resources through natural language, using an interactive conversational interface. In biomedical research, conversational AI has the potential to bridge the gap between human language and structured data, lowering barriers to complex knowledge systems and facilitating interdisciplinary access [20–22]. Unfortunately general-purpose chatbots such as ChatGPT (https://chatgpt.com/), Claude (https://claude.ai), and Gemini (https://gemini.google.com/) often struggle with domain-specific queries, source transparency, and hallucination, limiting their usability for scientific use [23–29]. These challenges underscore the need for systems that combine conversational flexibility of LLMs with direct integration into expert-curated knowledgebases to simultaneously achieve scientific rigor and user accessibility[30].

Here, we describe React-to-Me (https://reactome.org/chat), a domain-specific conversational interface designed to support intuitive, natural language exploration of the Reactome Pathway Knowledgebase. We describe the design of the system, objective benchmarking of response quality, and the results of user acceptance testing.

## Results

React-to-Me (Fig. 1) is a conversational interface that enables researchers to query the Reactome Pathway Knowledgebase in natural language and receive detailed and citation-linked responses grounded in curated pathway data. The system employs a Retrieval-Augmented Generation (RAG) framework to anchor an open-domain large language model (LLM) in curated Reactome data, ensuring factual precision, source traceability and biological relevance. Additional components provide guardrails against fictitious responses (“hallucinations”) and out-of-scope user questions by enforcing grounding and safety constraints at each stage of interaction. Although React-to-Me has been optimized for deployment with OpenAI’s GPT-4o-mini, its architecture is model-agnostic and compatible with any open-source or locally hosted LLM backend, supporting secure and adaptable deployment across diverse research environments.

**Figure 1.**
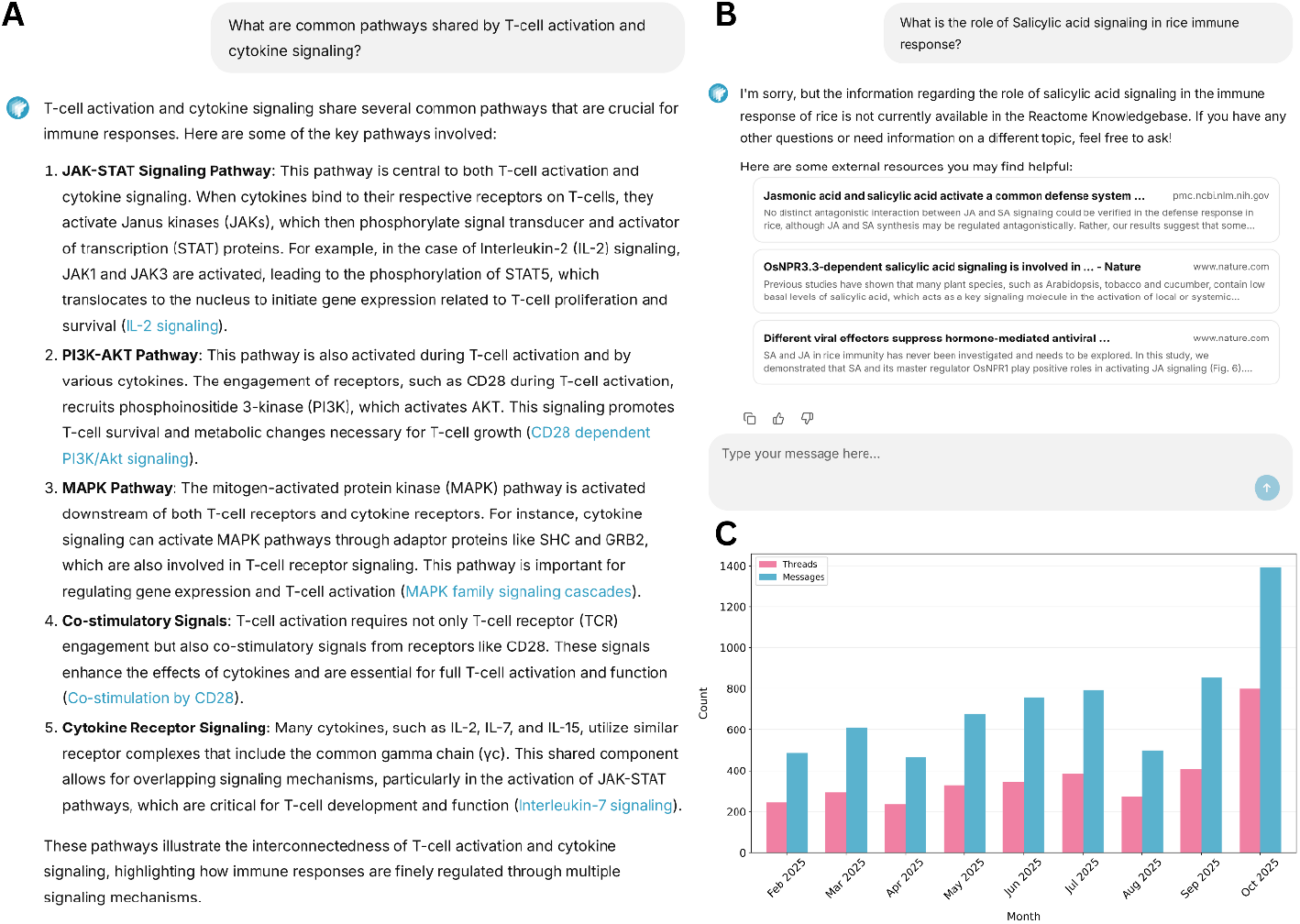
Overview of context-aware query handling by the React-to-Me conversational interface. **(A) Representative screen capture of system response to an in-scope biological query**. React-to-Me was queried about pathways shared by T-cell activation and cytokine signaling. A mechanistically detailed response is returned with direct citations to curated Reactome entries (blue hyperlinks), summarizing five key pathways (JAK-STAT, PI3K-AKT, MAPK, co-stimulatory signaling, cytokine receptor signaling). Each assertion is directly linked to supporting Reactome entries, enabling verification and direct navigation to curated pathway, reaction and entity records. **(B) Representative screen capture of system response to an out-of-scope (Plant biology) query**. For queries outside Reactome’s coverage (salicylic acid signaling in rice), React-to-Me refrains from speculative generation, explicitly indicates knowledge limitations, and instead provides links to relevant primary literature and external trusted biomedical resources for user-directed exploration. **(C) User engagement following public release**. Combined unauthenticated and authenticated user activity from February through October 2025, showing conversation threads initiated (pink) and total messages exchanged (blue).

### Architecture

React-to-Me employs a modular, multi-stage architecture designed to ensure scientific accuracy, source transparency, domain specificity and contextual coherence (Fig. 2A). The system processes each user query through five sequential components that together enforce strong grounding in Reactome’s curated content and prevent speculative or unsupported outputs.

**Figure 2.**
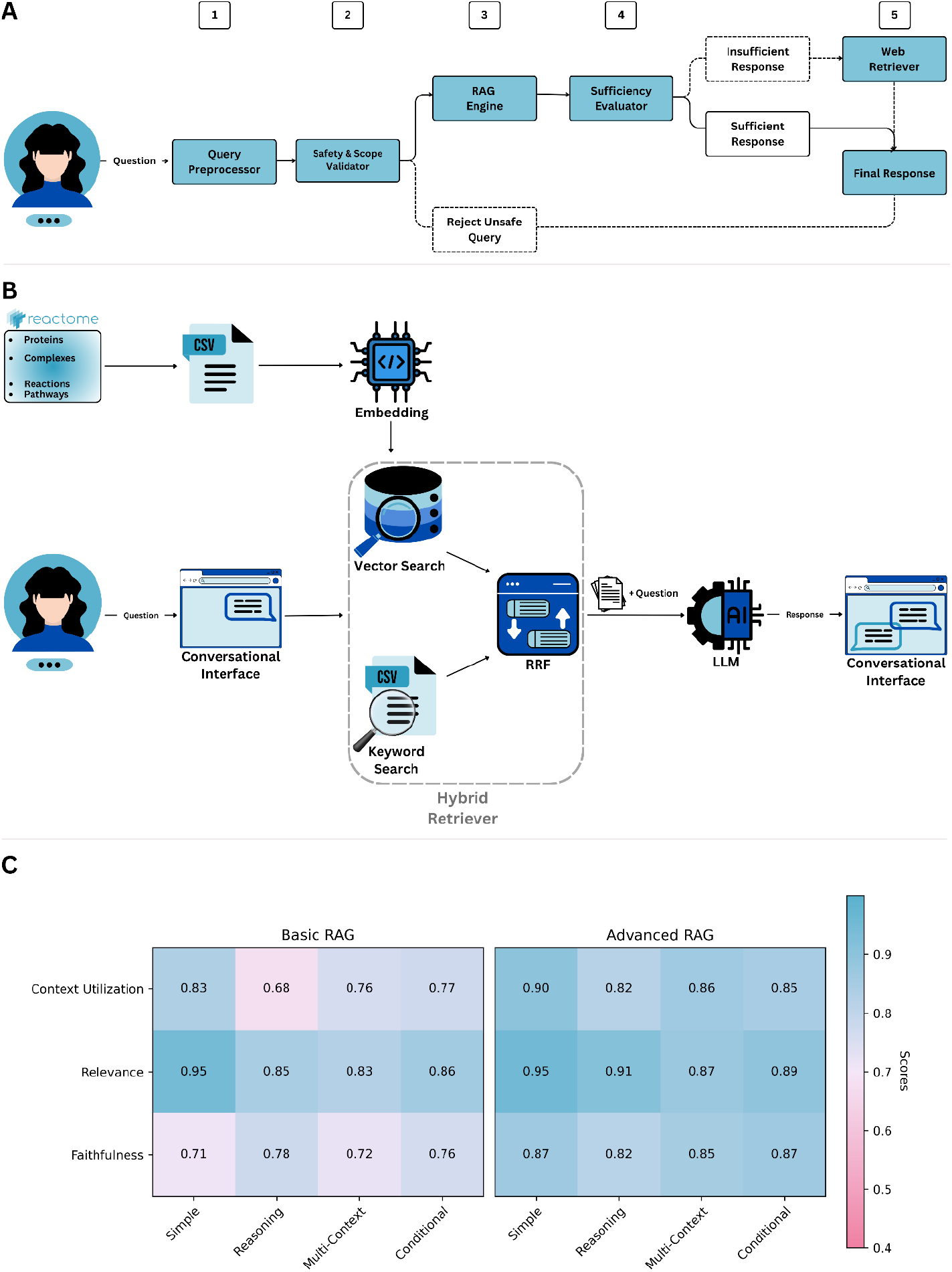
System architecture and benchmarking of the React-to-Me conversational AI framework. **(A) Modular pipeline for grounded biomedical question answering**. The React-to-me workflow consists of 5 sequential modules: query preprocessing, safety and scope validation, retrieval-augmented generation (RAG), response sufficiency evaluation, and fallback external retrieval. **(B) Hybrid retrieval engine integrating dense and sparse search**. Reactome pathway, reaction, and molecular entity records are embedded for semantic vector search and concurrently indexed for keyword-based retrieval. Results from both retrieval modalities are merged using reciprocal rank fusion (RRF) for balanced precision and recall. Aggregated context is supplied to the LLM for grounded response generation. **(C) Quantitative benchmarking of retrieval strategies using the Ragas evaluation framework**. Performance of a dense-only vector search baseline (Basic RAG) and the hybrid RRF-based configuration (Advanced RAG) was evaluated across three metrics (context utilization, relevance, and faithfulness) on 1,000 biomedical queries spanning four task types (simple, reasoning, multi-context, and conditional). Scores were normalized to [0, 1], with higher values indicating better performance. Advanced RAG showed consistent gains across all metrics, indicating improved contextual grounding and factual reliability.

#### Query Preprocessor

The system reformulates each user question into a standardized and precise query optimized for downstream retrieval components. This module normalizes biological terminology, resolves potentially ambiguous gene/protein names and pathway identifiers, and incorporates relevant conversational history to avoid semantic drift. The resulting clarified prompt ensures that the system interprets user intent consistently across multi-turn exchanges and grounding fidelity **(see Supplementary Methods §1.2.2)**.

#### Scope Validator

Before retrieval, each refined query is evaluated by an LLM-based classifier for domain relevance, and compliance with scientific and ethical guidelines **(see Supplementary Methods §1.2.3)**. Queries deemed unsafe, biologically irrelevant or otherwise inappropriate are filtered out and flagged. Prompt rejection at this stage preserves the system’s scientific integrity and ensures that only valid biological questions progress through the RAG pipeline.

#### RAG Module

Validated queries are processed through a RAG-Fusion pipeline that forms the core of React-to-Me’s grounding mechanism **(see Supplementary Methods §1.2.4 & 1.2.5)** [31]. Retrieval is performed over a curated index of Reactome pathways, reactions, molecular entities and associated annotations using two complementary strategies: (i) dense semantic retrieval, in which semantic vector embeddings capture contextual similarity among pathway descriptions, reactions, and molecular interactions; and (ii) sparse lexical retrieval, using BM25-based keyword matching to preserve sensitivity to exact biological identifiers (e.g., gene symbols, protein names) [32]. Results from these retrieval modes are integrated and re-ranked via Reciprocal Rank Fusion (RRF) to balance precision and recall (Fig. 2B) [33]. The top-ranked entries are concatenated and injected into a structured contextual prompt that is passed to a prompt-constrained, near-deterministic LLM (temperature = 0.0). This generation stage synthesizes a coherent natural-language response strictly while strictly adhering to retrieved Reactome evidence. Each factual assertion is hyperlinked to its corresponding Reactome entry, ensuring transparency, traceability, and alignment with curated biological knowledge. This design suppresses the LLM’s tendency to engage in speculative generation and maintains factual alignment across responses.

#### Sufficiency Evaluator and Fallback Retrieval

After answer generation, an LLM-based sufficiency evaluator determines whether the generated response adequately addresses the user’s question **(see Supplementary Methods §1.2.6 & 1.2.7)**. When coverage is incomplete, such as for emerging biological mechanisms, rare interactions or pathways in non-human species, React-to-Me does not generate speculative context. Instead, the system initiates a fallback retrieval module to identify and provide users with links to external relevant and trusted biomedical repositories (e.g. PubMed Central, NIH databases) to guide further user-directed exploration (Fig. 1B). This strict separation between grounded generation and exploratory retrieval reflects a conservative design principle aimed at maintaining factual integrity.

### Computational Benchmarking of Knowledge Retrieval Strategies

The retrieval module is the most critical determinant of overall system performance, as it establishes the completeness, precision and biological specificity of the information provided to the language model. Insufficient retrieval restricts the model’s ability to construct comprehensive answers, while overly broad or irrelevant context can dilute biological specificity and reduce factual precision. To quantify the impact of retrieval design on output quality, we systematically compared two alternative retrieval configurations using the Retrieval-Augmented Generation Assessment (Ragas) evaluation framework [34].

The baseline configuration (“Basic RAG”) employed dense retrieval over embedded Reactome entries. The enhanced configuration (“Advanced RAG”) incorporated both dense and sparse lexical retrieval, integrating their rankings through Reciprocal Rank Fusion (RRF) prior to generation (Fig. 2B). We hypothesized that this hybrid strategy would improve factual grounding by combining the contextual flexibility of semantic matching with the precision of a keyword-based search.

Evaluation was performed on a synthetic dataset of 1,000 biologically grounded questions derived from Reactome content using the Ragas test set generation workflow **(see Supplementary Methods §1.3)**. The dataset spanned four representative query types: simple factual lookups (Simple), mechanistic reasoning (Reasoning), multi-context synthesis (Multi-context), and conditional logic (Conditional) [34]. Each system was assessed across three core Ragas metrics: context utilization (extent to which retrieved evidence was incorporated into the response), relevance (semantic alignment with the query), and faithfulness (factual consistency with the retrieved context).

The Advanced RAG configuration consistently outperformed the Basic RAG baseline across all evaluation dimensions (Fig. 2C). Notably, relevance scores remained comparably high in both configurations, suggesting that dense retrieval alone is often sufficient for retrieving semantically appropriate context. However, the Basic RAG system showed consistent deficits in faithfulness, indicating that semantic similarity alone does not reliably guarantee factual alignment. In contrast, the hybrid retrieval strategy substantially improved both context utilization and faithfulness, yielding responses that were richer in mechanistic detail, more accurately grounded in Reactome data and less prone to unfounded assertions. These results establish the retrieval architecture as a primary determinant of factual reliability in domain-specific RAG systems.

### Expert Evaluation of Grounded Vs. Open-domain Responses

To determine whether grounding improves expert-perceived quality of biological explanations, we conducted a structured, blinded expert evaluation comparing React-to-Me to an ungrounded baseline LLM (GPT-4o-mini). Ten blinded domain experts independently evaluated paired responses to 15 molecular biology questions stratified to two complexity categories: query-like tasks requiring factual recall and definitional understanding, and reasoning tasks requiring synthesis and causal or multi-step inference. Each response was scored using a standardized four-point rubric across three metrics: factual accuracy, level of granularity, and relational depth **(see Supplementary Methods §1.4)**. Inter-rater reliability for absolute scores on shared questions was low (ICC(2,1) range: -0.05 to 0.18), consistent with the inherent subjectivity of the task. Accordingly, statistical analyses employed cumulative-link mixed-effects models with random intercepts for participants and questions, as well as system-specific random slopes to account for individual calibration differences and repeated-measures structure **(see Supplementary Methods §1.4.3)**. This framework ensures that estimated system-level contrasts reflect the true performance differences rather than rater-specific variability. Robustness was confirmed using nonparametric Wilcoxon and sign tests (Supplementary Table S2-S3).

#### Grounding yields significant system wide performance gains

Across all evaluators, questions and scoring dimensions, React-to-Me consistently outperformed the ungrounded baseline (Table 1; Table S1). Mixed-effects ordinal regression indicated that grounded responses were twice as likely to receive higher scores (OR = 2.01 [95 % CI: 1.50-2.69], *Holm-adj p < 0*.*01*; Table 1), corresponding to a mean gain of +0.26 points on the four-point scale (*Wilcoxon p < 0*.*01*; Table 1), and a 60% probability that any randomly selected grounded response would surpass its ungrounded counterpart (CLES = 0.60 ± 0.03; Table 1). Nonparametric analyses confirmed these findings. Grounded responses were preferred in 71.6% of paired comparisons (154 vs. 61; 235 ties ; p < 0.01; Table S2), and all ten evaluators showed an aggregate preference for React-to-Me overall ( *p < 0*.*01*; Table S3). Estimated statistical power exceeds 0.99 across all comparisons (Table 1; Table S2).

**Table 1.** Statistical comparison of expert ratings for grounded versus ungrounded responses. Cumulative-link mixed-effects model results comparing React-to-Me (grounded) to GPT-4o-mini baseline (ungrounded) across 450 paired evaluations (10 experts × 15 questions × 3 metrics). OR: odds ratio for receiving a higher rating category; Holm-adj p: family-wise error corrected p-value; CLES: common language effect size (probability grounded response scores higher); Mean Δ: average score difference in points on 4-point scale. Models included random intercepts for evaluator and question, with system-specific random slopes. All comparisons favor React-to-Me (OR > 1, p < 0.05).

| Stratum | N (pairs) | OR [95% CI] | Holm-adj $p$ | Wilcoxon $p$ | CLES (inclusive) | Mean $\Delta$ (pts) | Power |
| --- | --- | --- | --- | --- | --- | --- | --- |
| Overall | 450 | 2.01 [1.50–2.69] | $2.9 \times 10^{-6}$ | $1.7 \times 10^{-11}$ | 0.60 (0.57–0.63) | +0.26 | 1.000 |
| Query | 270 | 1.89 [1.34–2.66] | $5.7 \times 10^{-4}$ | $7.1 \times 10^{-7}$ | 0.60 (0.56–0.63) | +0.24 | 1.000 |
| Reasoning | 180 | 2.20 [1.17–4.16] | 0.015 | $2.5 \times 10^{-6}$ | 0.61 (0.57–0.66) | +0.29 | 0.999 |
| Factual accuracy | 150 | 2.97 [1.68–5.24] | $5.3 \times 10^{-4}$ | $2.0 \times 10^{-6}$ | 0.61 (0.57–0.65) | +0.26 | 1.000 |
| Level of granularity | 150 | 2.88 [1.62–5.11] | $6.4 \times 10^{-4}$ | $1.2 \times 10^{-5}$ | 0.63 (0.57–0.68) | +0.33 | 0.998 |
| Relational depth | 150 | 1.83 [1.08–3.09] | 0.023 | $2.3 \times 10^{-3}$ | 0.58 (0.52–0.63) | +0.19 | 0.893 |

The largest and most consistent performance gains were observed in factual accuracy (Fig. 3A). Grounded responses were nearly three times more likely to receive higher ratings (OR = 2.97 [1.68-5.24]; *Holm-adj p < 0*.*01*; Table 1), outperforming the baseline in 82% of paired comparisons (p < 0.01; Table S2).

**Figure 3.**
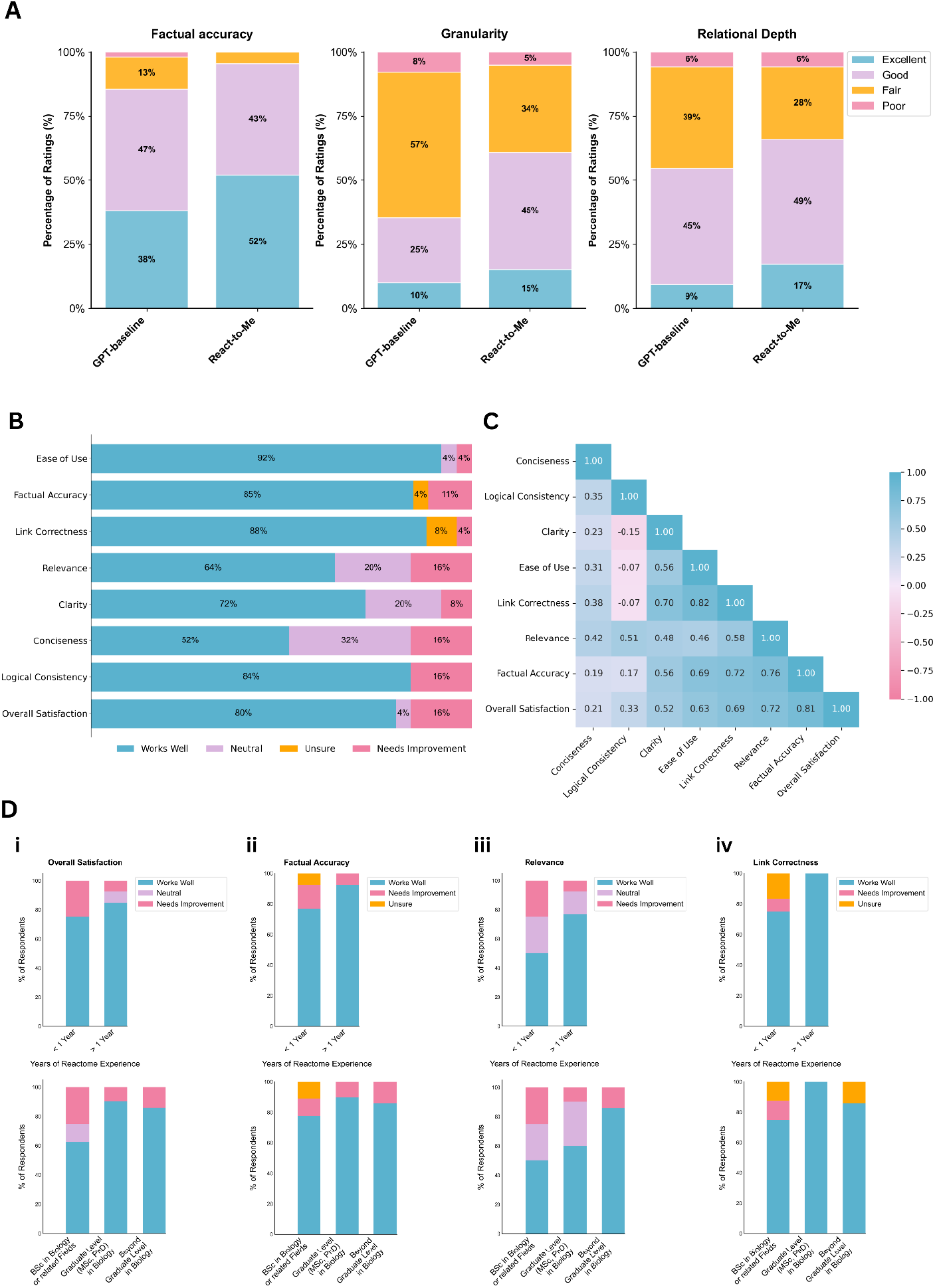
User evaluation of React-to-Me and perceived quality. **(A) Expert rating distributions for React-to-Me versus GPT-4o-mini baseline across three evaluation metrics**. React-to-Me showed higher proportions of “Excellent” ratings for factual accuracy (52% vs 38%), granularity (15% vs 10%), and relational depth (17% vs 9%) (n=150 paired evaluations: 10 experts × 15 questions each, 4-point scale). **(B) Aggregated user ratings across eight performance metrics**. Results from the user satisfaction survey (n=27) assessing ease of use, factual accuracy, link correctness, relevance, clarity, conciseness, logical consistency and overall satisfaction. Most respondents rated performance positively across all dimensions, with the highest approval for ease of use, link correctness and factual accuracy. **(C) Pairwise Pearson correlation matrix of evaluation metrics**. Pearson correlation coefficients highlight strong positive relationships between factual accuracy, relevance and overall satisfaction, demonstrating that factual reliability and source transparency are the primary determinants of user trust. Correlation coefficients are shown with color intensity (blue: positive, pink: negative). **(D) Stratified analysis by user domain expertise**. Top row: stratification by Reactome experience (<1 year vs ≥1 year); bottom row: stratification by education level in biology. Each panel shows four key metrics with stacked bars indicating rating distributions. Higher expertise is consistently associated with higher satisfaction ratings.

Similarly, grounding substantially improved biological specificity and granularity. React-to-Me responses showed 2.9-fold higher odds of superior granularity ratings (OR = 2.88 [1.62-5.11]; *Holm-adj p* < 0.01; Table 1), with superiority observed in 71% of paired comparisons (*p* < 0.01; Table S2). Although the participant-level preference was not significant (8 of 10 raters, p = 0.0547; Table S3) grounded responses showed the largest median score gain among all metrics (Mean Δ = +0.33; Wilcoxon *p* < 0.01; Table 1).

Improvements in relational depth were more moderate but remained significant. Grounded responses exhibited 1.83-fold higher odds of superior ratings (OR = 1.83 [1.08-3.09]; *Holm-adj p* < 0.05; Table 1), outperforming the baseline in 65% of paired evaluations (p < 0.01; Table S2).

Performance gains were consistent across both factual recall and reasoning-oriented tasks, indicating that its benefits extend beyond knowledge retrieval to higher-level biological reasoning. Grounded answers were nearly twice as likely to receive higher expert ratings on query-like questions (OR = 1.89 [1.34-2.66] ; p < 0.01; Table 1), and more than twice as likely on reasoning questions (OR = 2.20 [1.17-4.16]; p < 0.05; Table 1). Mean score gains of +0.24 and +0.29 points respectively, suggested slightly greater improvements for tasks requiring synthesis and mechanistic reasoning. Expert preferences mirrored these model-based effects. Nine out of ten evaluators favored React-to-Me for both query and reasoning tasks (p < 0.05 and p < 0.01 respectively; Table S3). Grounded responses outperformed the ungrounded baseline in 71% of factual recall and 72.5% of reasoning tasks (p < 0.01; Table S2). Together, these results show that grounding consistently enhances expert-perceived response quality across question types, strengthening both factual precision and inferential coherence required for causal and multi-step reasoning.

### User Satisfaction

To evaluate the broader usability and perceived quality of React-to-Me for community users of the Reactome Knowledgebase, we deployed an online survey assessing the chatbot across eight key performance metrics: Ease of Use, Clarity, Conciseness, Logical Consistency, Link Correctness, Relevance, Factual Accuracy and Overall Satisfaction. Participant recruitment was conducted through local institutional bulletins and global online outreach via trusted scientific communities and networks. Over a 29-week period we received responses from 27 anonymous participants representing a diverse cohort of biomedical professionals, educators and students who provided structured ratings and optional qualitative feedback on the system’s performance (Fig. S1A-B).

Overall, survey results indicate strong user satisfaction across key dimensions (Fig. 3A). The highest ratings were observed for ease of use (92%), link correctness (88%) and factual accuracy (85%), suggesting that users found the interface intuitive, the citations reliable and the responses scientifically trustworthy. Although ratings for conciseness, logical consistency and relevance were generally favourable, a minority of respondents identified areas for improvement.

To identify features that most strongly influence user satisfaction, we conducted a Pearson pairwise correlation analysis across all evaluation metrics (Fig. 3B). Factual accuracy emerged as the strongest predictor of overall satisfaction (*ρ* (rho) =0.81), followed by relevance (*ρ* (rho) =0.72), link correctness (*ρ* (rho) =0.69) and ease of use (*ρ* (rho) =0.63). In contrast, stylistic attributes such as conciseness (*ρ* (rho) =0.21) and logical consistency (*ρ* (rho) =0.33) were only weakly associated with satisfaction, indicating that while linguistic fluency contributes to perceived quality, users ultimately prioritize factual reliability, contextual relevance and source transparency.

To investigate how domain familiarity shapes user experience we conducted a stratified analysis of survey responses based on self-reported biology education level and Reactome usage experience (Fig. 3D, S1C). Although subgroup analyses were underpowered due to modest sample sizes, descriptive trends indicate that participants with more advanced biological training gave higher “works well” ratings for several evaluation metrics. Differences were most pronounced for Relevance (BSc: 50.0%; Graduate: 60.0%; Beyond Graduate: 85.7%; Fig. 3D(iii)), Overall Satisfaction (BSc: 62.5%; Graduate: 90.0%; Beyond Graduate: 85.7% Fig. 3D(i)), and Ease of Use (BSc: 75.0%; Graduate/Beyond Graduate: 100.0%) (Fig. S1-C(i)).

Similar patterns were observed when stratified by Reactome experience. Users with ≥ 1 year of Reactome familiarity rated the system more favourably for relevance (+26.9 percentage points; 3D(iii)), clarity (+26.3 pp; Fig. S1-C(ii)) and link correctness (+25.0 pp, reaching 100.0% “works well”; Fig. 3D(iv)). In contrast, stylistic attributes such as conciseness and logical consistency showed weaker and less consistent variation with expertise. Overall, these subgroup analyses point to possible expertise-related differences in perceived relevance, factual reliability and usability.

## Discussion

This work introduces React-to-Me, a domain-specific conversational assistant that provides intuitive, natural-language access to complex biological knowledge from the Reactome Knowledgebase. By grounding LLM generation in retrieved pathway records, React-to-Me delivers detailed and factually accurate responses that maintain the source traceability required for biomedical research applications.

Across computational benchmarking, expert evaluation, and community user satisfaction surveys, React-to-Me consistently demonstrated superior performance relative to baselines. Systematic comparison of retrieval architectures using the Ragas framework confirmed that hybrid dense-sparse retrieval with reciprocal rank fusion substantially improved both context utilization and faithfulness, yielding responses more accurately aligned with Reactome data and less prone to hallucination. Expert evaluation further confirmed that grounding substantially enhanced factual accuracy, biological granularity and mechanistic depth. User satisfaction surveys corroborated these findings, reporting high satisfaction with ease of use, citation reliability, and scientific trustworthiness. Factual accuracy emerged as the strongest predictor of overall satisfaction (ρ (rho) = 0.81), indicating that users prioritize verifiable, source-grounded information over linguistic fluency.

These findings demonstrate that React-to-Me enables intuitive exploration and engagement with Reactome’s curated knowledge, while preserving the accuracy and contextual specificity essential for research applications. By enabling natural language interaction with complex pathway data, React-to-Me facilitates hypothesis generation, exploration of pathway interactions, and provides an opportunity to develop expert LLM-assisted research workflows.

A central conclusion of this study is that grounding an LLM in curated biological knowledge confers substantial benefits across both factual recall and reasoning-oriented tasks, suggesting that RAG serves as more than a simple lookup mechanism. A plausible interpretation is that access to biologically validated relationships among molecules, reactions, and regulatory events, constrains the LLM to construct explanations along biologically plausible and causally coherent trajectories. In contrast, ungrounded models operating purely on statistical co-occurrence may more readily generate mechanistically inconsistent explanations, particularly in complex or sparsely represented areas of biology.

Beyond enabling conversational access to Reactome content, the LLM-powered architecture introduces several additional benefits for the React-to-Me system. Multilingual capabilities lower accessibility barriers for non-English speaking users and broaden the reach of Reactome beyond its traditional user base. The system’s ability to adapt explanations to user intent and expertise extends its utility to clinicians, educators, students, and the broader public. Moreover, the evidence-aware fallback mechanism maintains transparency at the boundaries of Reactome’s coverage by transparently acknowledging gaps and directing users to primary literature or other external authoritative sources without resorting to speculative generation.

Despite these strengths, it is important to acknowledge several limitations in the current system. In order to prioritize factual accuracy, React-to-Me’s scope is deliberately constrained to the human biological pathways currently curated in Reactome, which limits coverage of model organism biology, comparative analyses, and cross-species pathway mapping. The system’s reliance on expert-curated knowledge also means that coverage inevitably lags behind rapidly emerging biological mechanisms, rare molecular interactions, and newly discovered pathway components that have not yet undergone expert curation. Although strict grounding, conservative prompting and rigorous safeguards substantially reduce the risk of hallucinations, the probabilistic nature of language model generation means that factually inconsistent outputs remain possible, particularly for queries at the boundaries of Reactome’s curated scope or when biological terminology is ambiguous. In addition, the system’s current deployment uses a commercial language model API (GPT-4o-mini), raising considerations around long-term cost, data governance, and long-term sustainability for institutional deployments. Although the architecture is model-agonistic, practical migration to open-source or locally hosted models will depend on continued advances in domain performance and infrastructure support.

While the general user survey indicated that overall user satisfaction was high, respondents’ feedback identified several areas for improvement. Although clarity, conciseness and logical consistency were generally rated positively, they also emerged as opportunities for further enhancement. These observations suggest that even when factual reliability is high, improvements in linguistic coherence could further enhance usability and response quality.

Future development will focus on expanding both the breadth and depth of grounded knowledge. Integration with additional curated biological resources (UniProt, Alliance of Genome Resources) via the Model Context Protocol (MCP) will extend coverage of molecular entities and cross-species pathways [35–40]. Ongoing efforts aim to refine interaction and accessibility for clinical researchers, undergraduate students, and informed lay users while preserving precision and verifiability for expert applications. As React-to-Me continues to evolve through community engagement and technical refinement, it will serve as a concrete example of how domain-specific conversational AI can democratize access to specialized biological knowledge without sacrificing the accuracy and transparency that underpin scientific trust.

## Data availability

The React-to-Me conversational interface is freely accessible at https://reactome.org/chat and will be maintained at this URL for a minimum of five years following publication. Source code for the system implementation, including deployment instructions, is available under the MIT license at https://github.com/reactome/reactome_chatbot. Statistical analysis scripts (R) for all computational analyses and figure generation are available in the /analysis directory of the GitHub repository.

Expert evaluation data, including the 15 molecular biology questions, 450 paired response ratings, and the original React-to-Me and GPT-4o-mini responses assessed by curators, are publicly available on Zenodo (DOI: 10.5281/zenodo.17602575). The Reactome pathway data used for retrieval is publicly available at https://reactome.org/download-data.

## Acknowledgements

We thank the Reactome team for their contributions to system development, including expert evaluation of responses and valuable feedback throughout the design and implementation process: Peter D’Eustachio, Marc Gillespie, Bruce May, Karen Rothfels, Ralf Stephan, Lisa Matthews, Veronica Shamovsky. We are grateful to the external molecular biology experts who participated in the blinded comparative evaluation, and to the graduate researchers and biomedical professionals who completed the user satisfaction survey.

