## Supplementary information for "React-to-Me: A Conversational Interface for Interactive Exploration of the Reactome Pathway Knowledgebase"

---

### Table of Contents


Supplementary Methods 3

1 Retrieval-Augmented Generation (RAG) Framework 4

2 React-to-Me System Implementation. 4

2.1 Corpus Construction and Embedding 4

2.2 Query preprocessing and standardization 4

2.3 Safety and scope validation 5

2.4 Hybrid semantic-lexical retrieval 5

2.6 Evidence sufficiency evaluation 6

3 Computational Benchmarking of Retrieval Strategy Using Ragas 7

4 Expert Evaluation 8

5 General User Satisfaction Survey 13

References 18

### Supplementary Methods

The React-to-Me chatbot was developed as a modular, containerized Python application designed for reproducibility, extensibility, and secure deployment in biomedical research contexts. It supports Python 3.12, and is maintained on GitHub (https://github.com/reactome/reactome_chatbot). Software dependencies are explicitly specified and version-locked using Poetry, with continuous integration workflows ensuring environment consistency across supported versions.

The system architecture comprises multiple coordinated components:

- Frontend interface: Built using Chainlit (v2.0) (https://docs.chainlit.io/get-started/overview), served with HTTPS via NGINX for secure web access.
- Backend orchestration: Managed by LangGraph (v0.2) (https://www.langchain.com/langgraph), which coordinates a multi-agent workflow for query interpretation, retrieval, generation, and post-processing.
- Retrieval and generation modules: Implemented using LangChain (v0.3) (https://www.langchain.com/), with hybrid retrieval via rank-bm25 (v0.2.2) and dense similarity vector search with ChromaDB (v0.5) (https://www.trychroma.com/) using HNSW indexing. Language generation is performed using GPT-4o-mini (https://platform.openai.com/docs/models/gpt-4o-mini).
- Embedding generation: Reactome entries are individually indexed using LangChain's CSV loader and embedded using OpenAI's text-embedding-3-large model (https://platform.openai.com/docs/models/text-embedding-3-large). Reactome entries are individually indexed and embedded using LangChain's CSV loader.

React-to-Me supports two access modes: an unauthenticated guest mode, which enables ephemeral, anonymous interaction without storing any conversation history, and a registered login mode, which preserves session history to support multi-turn reasoning and longitudinal use. No personally identifiable information (PII) is collected in either mode, and all usage data from the login mode is securely stored in compliance with ethical data governance standards.

The project is distributed under the permissive Apache License 2.0 to encourage downstream reuse and collaborative development.

### 1 Retrieval-Augmented Generation (RAG) Framework

To enable natural language interaction with Reactome's structured pathway knowledge, we developed React-to-Me using state-of-the-art Retrieval-Augmented Generation (RAG) techniques [1–4]. RAG systems combine the capabilities of large language modes (LLMs) with real-time retrieval of relevant information from trusted knowledge sources, ensuring that answer generation is anchored in retrieved evidence rather than the model's pretrained internal knowledge. This approach mitigates the key limitations of open-domain chatbots and fine-tuning approaches which internalize knowledge within model weights, and cannot guarantee alignment with source material [5–9]. For scientific applications where factual precision and contextual relevance are essential, RAG provides a principled mechanism for ensuring that generated responses remain anchored in Reactome's expert-curated content [10,11].

The React-to-Me implements a modular RAG pipeline tailored to Reactome's ontology and content structure. The workflow includes three coordinated stages: (1) hybrid retrieval from a preprocessed Reactome corpus, (2) context-aware prompt construction and generation, and (3) post-hoc answer validation, including fallback retrieval for ambiguous queries. This modular architecture enables independent optimization of each pipeline component and maintains full traceability from generated responses back to specific Reactome pathway entries.

### 2 React-to-Me System Implementation.

##### 2.1 Corpus Construction and Embedding

The backend ingests the most recent React-to-Me release in CSV format. Each record is processed using LangChain's CSVLoader together with a custom preprocessing pipeline (metadata_csv_loader.py) that extracts and concatenates four metadata fields: st_id, display_name, canonical_geneName, and synonyms_geneName. These are stored alongside cleaned text content as individual atomic documents.

All documents are embedded using OpenAI text-embedding-3-large model, yielding 3072-dimensional vectors that capture semantic similarity between Reactome entries. Embeddings are stored in ChromaDB (v0.5) with HNSW indexing for efficient approximate nearest-neighbor search, accompanied by JSON metadata including Reactome ID, entity type and canonical URLs. Index construction is updated periodically through a dedicated build script (embeddings_manager), but auto-refresh is disabled in production to ensure stability and reproducibility across deployments.

##### 2.2 Query preprocessing and standardization

To ensure that downstream components operate on well-formed, domain-specific and context-aware questions, prior to retrieval, each user question is reformulated using a prompt-engineered LLM preprocessor (Fig. 2A). The prompt instructs the model to:

- Resolve co-references and contextual dependencies from multi-turn conversations.
- Normalize biological terminology (e.g., gene/protein names, pathway identifiers).

- Correct grammar and reduce linguistic ambiguity.
- Optimize phrasing for both semantic vector search and case-sensitive lexical keyword-based matching.

If the user's query is already well-formed and self-contained, it is returned unchanged. The preprocessor never supplies additional biological facts or answers the question; the sole purpose is to standardize the user inputs for retrieval.

**2.3 Safety and scope validation**

Prior to retrieval, every refined query is screened by an LLM-based safety and scope classifier (Fig. 2A). This classifier operates using a structured prompt and evaluates the query according to the following criteria:

- Domain relevance: the query must pertain to biology, molecular mechanisms or related life-science topics.
- Appropriateness: the query must not contain offensive, discriminatory or otherwise inappropriate content
- Safety: the query must not request or imply medical advice, harmful, dual-use or unethical biological activities. Ambiguous or potentially harmful queries are treated as unsafe.

Queries are returned as "Safety: "true" or "false" with a brief justification when unsafe. Flagged queries are blocked and do not proceed to retrieval or generation. This early gating mechanism aligns the system with best practices for responsible AI deployment in biological contexts.

**2.4 Hybrid semantic-lexical retrieval**

Validated queries are processed through React-to-Me's hybrid retrieval engine, which integrates dense semantic search with sparse lexical marching to maximize coverage across Reactome's heterogeneous ontology (Fig. 2B).

Dense semantic retrieval: Textual content from Reactome is embedded according to Supplementary Methods 1.2.1. Cosine similarity search over the embeddings stored in ChromaDB identifies semantically relevant entries, capturing relationships embedded in pathway and reaction descriptions and molecular function annotations.

Sparse lexical retrieval: In parallel, all documents are indexed using BM25 (rank-bm25, v0.2.2). This retrieval strategy enables precise matching of gene symbols, protein names, and pathway identifiers that may not be well-represented in embedding space [12].

Reciprocal rank fusion (RRF): results from the dense and sparse retrieval modes are integrated using Reciprocal Rank Fusion (RRF), which promotes documents identified by both strategies while preserving complementary hits [13,14]. The ranked list is truncated to the top 60 documents, a threshold selected to balance evidence breadth against LLM context-window limits.

##### 2.5 Prompt construction and grounded answer generation

Grounded response generation is performed using GPT-4o-mini, operated deterministically with temperature = 0.0 to ensure reproducibility across repeated queries. Responses are streamed to maintain conversation fluidity.

React-to-Me uses a structured prompt template with the following components:

- Role and scope definition: the model is instructed to act as an expert molecular biologist with access strictly limited to the provided Reactome context.
- Grounding and prohibition rules: the prompt explicitly forbids speculation or use of external knowledge and instructs the model to acknowledge when Reactome lacks relevant information.
- Citation requirements: the model must cite every factual statement using standardized reactome hyperlinks.
- Accessibility constraints: responses must be clear, concise and suitable for researchers, educators and clinicians.

Retrieved documents are concatenated with the refined query and injected into this template. If the context window is exceeded, lower-ranked items (see Supplementary Methods 1.2.4) are omitted to preserve high-value evidence. If no documents support the question, the model is instructed to decline to answer and defer to the fallback mechanism.

##### 2.6 Evidence sufficiency evaluation

After answer generation, a dedicated LLM-based sufficiency evaluator determines whether the response fully addresses the user's question. Using a concise binary grading prompt ("Yes"/"No"), the evaluator assesses:

- Completeness of the biological explanation.
- Adequacy of contextual background.
- Fidelity to the user's intent.

If the response is deemed complete, it is returned to the user. If incomplete, the fallback retrieval workflow is initiated.

##### 2.7 Evidence-aware fallback retrieval

For queries that exceed Reactome's coverage, such as emerging pathways, rare interactions, or non-human processes, React-to-Me employs a structured fallback mechanism that avoids speculative generation while still supporting user inquiry (Fig. 1B). The fallback workflow is orchestrated via LangGraph and proceeds as follows:

- Query reformulation: the original user query is refined to optimize compatibility with external search.

- External retrieval: the refined query is submitted to the Tavily search API (https://www.tavily.com/), configured to prioritize high-quality biomedical repositories including PubMed Central (https://pmc.ncbi.nlm.nih.gov/), NIH databases (https://www.ncbi.nlm.nih.gov/) and related sources .
- Transparent handoff: Importantly, React-to-Me does not attempt to use the retrieved content for further generation. Instead the system informs the user that Reactome does not contain relevant information and returns a list of external sources with direct links to guide further user directed exploration.

This design enforces a strict separation between curated knowledge generation and external retrieval, preserving factual integrity and user trust. By avoiding speculative generation and clearly highlighting the limits of its internal knowledge, React-to-Me maintains transparency and reliability even when definitive answers cannot be internally generated.

#### 3 Computational Benchmarking of Retrieval Strategy Using Ragas

To assess the impact of retrieval architecture on answer quality, we benchmarked two versions of the React-to-Me system that differed only in their retrieval strategy. The 'Basic RAG' configuration relied solely on dense semantic retrieval over embedded Reactome entries, whereas the 'Advanced RAG' configuration combined dense retrieval with sparse keyword-based search followed by Reciprocal Rank Fusion (RRF) (Fig. 2B). All other system components, including the query preprocessor, prompt templates, and GPT-4o-mini generation module were held constant to isolate the effects of retrieval strategy.

Benchmarking was conducted using the Retrieval-Augmented Generation Assessment (Ragas) framework, a task-agnostic, reference-free evaluation method designed to evaluate the quality of responses generated by retrieval-augmented systems [15]. Unlike traditional reference-based metrics such as ROUGE or BLEU, Ragas assesses the relationship between the retrieved context, original query, and the generated response, making it well suited for open-ended, knowledge-intensive tasks such as biomedical question answering [16,17].

A benchmarking dataset of 1,000 natural language queries were constructed based on Reactome content, spanning a range of biomedical question types: factual lookups, causal and mechanistic reasoning, multi-context synthesis, and conditionally scoped prompts. Each query was independently answered by both the Basic and Advanced RAG configurations, generating two model outputs per item. Ragas then automatically scored responses across three key dimensions:

- Context Utilization: The degree to which generated answers incorporated and reflected the retrieved evidence.
- Relevance: Semantic alignment between the response and the user's original intent.
- Faithfulness: Factual consistency of the answer relative to the retrieved context.

This benchmarking protocol provided a rigorous, context-aware method for evaluating retrieval strategies in a scientifically grounded question answering setting, and informed subsequent architectural decisions in the development of React-to-Me.

### 4 Expert Evaluation

#### 4.1 Survey Design

To compare grounded and ungrounded LLM responses in terms of factual reliability, biological specificity, and mechanistic depth, we conducted a structured, blinded expert evaluation using a curated set of 109 molecular biology questions derived Voet & Voet, Biochemistry (2nd ed.) [18]. The full question–answer dataset is publicly available on Zenodo (DOI: 10.5281/zenodo.17602575). Questions were stratified into two cognitive categories:

- Query-like (n = 76): tasks requiring factual recall and definitional understanding (e.g., “What is the complement system?”).
- Reasoning (n = 33): tasks requiring synthesis, causal reasoning, or multi-step inference (e.g., “Sublethal cyanide poisoning may be reversed by the administration of nitrites. These substances oxidize hemoglobin, which has a relatively low affinity for CN-, to methemoglobin, which has a relatively high affinity for CN-. Why is this treatment effective?”).

Ten domain experts participated in the study. Each participant evaluated 15 questions: five general biology questions common to all evaluators, and ten field-specific questions selected from a dropdown list to match their expertise. This ensured both consistency across participants and alignment with individual domain knowledge. Each evaluator reviewed 9 query-like and 6 reasoning questions, providing balanced coverage across the two cognitive categories..

For each question, curators assessed two LLM-generated responses:

- React-to-Me: grounded in Reactome data using hybrid RAG (see Supplementary Methods 1.2)
- Ungrounded general-purpose model: response generated by GPT-4o-mini, using the same prompt without external grounding.

To minimize potential knowledge gaps, a short background excerpt adapted from Voet & Voet accompanied each question [18–20]. These background hints were explicitly designated as optional context and not as scoring keys. Evaluators were instructed to evaluate the chatbot responses solely on their intrinsic scientific merit.

The identity of the chatbot system (React-to-Me vs. GPT-4o-mini) was fully blinded during evaluation. System names and source identifiers were removed from all materials. To minimize stylistic cues and standardize evaluation, both systems were prompted to generate responses of 150 words or fewer, and outputs were formatted to match each other’s overall presentation style. Presentation order of system responses was alternated on a per-question basis to mitigate potential order effects. This systematic alternation ensured balanced exposure and eliminated positional bias over the survey.

##### 4.2 Evaluation Criteria

Expert assessments were guided by a standardized evaluation toolkit designed to promote consistent scoring across participants and question types (Supplemental Appendix 1). Each response was evaluated along three predefined metrics:

- Factual Accuracy: Claims are factually correct, mechanistically sound, consistent with current scientific knowledge, and interpretable by both expert and non-expert readers.
- Level of Granularity: Precision and selectivity in identifying relevant biological entities (genes, proteins, pathways, reactions, diseases), with higher scores reflecting expert-level selectivity and integration.
- Relational Depth: Mechanistic and systems-level insight into how identified elements function and interact, capturing causal pathways, regulatory effects, and contextual relevance.

Each metric was scored on a four-point ordinal scale (1 = poor, 4 = excellent). Detailed descriptors, examples, and common pitfalls were provided for each level in the rubric. Curators were instructed to evaluate each response independently, without cross-comparing paired chatbot answers or textbook ‘optional hint’ reference excerpts. This evaluation framework ensured that judgments emphasized the intrinsic quality of each response, supporting fair, standardized comparison of domain-grounded and general-purpose chatbot outputs across both factual and reasoning-oriented tasks.

##### 4.3 Data Handling and Analysis

Survey responses were collected using Google Forms, exported to CSV, and stored securely on institutional servers. Source files were encrypted and deleted from the collection platform following transfer. No personally identifiable information (PII) was collected.

Each evaluation generated three scores per system, per question, yielding 900 total observations (10 participants, x 15 questions x 3 metrics x 2 systems) or 450 paired system comparisons. Scores were encoded as ordered factors (1 = poor, 4 = excellent), and the system identity was coded with ChatGPT as the reference condition. All analyses were conducted in R (v4.3.2) using the ordinal, effsize, boot, and pwr packages.

**Statistical Framework.** Analyses followed a prespecified hierarchical framework testing global and stratified hypotheses of system superiority while respecting the ordinal and repeated-measures structure of the data.

Primary inference employed cumulative-link mixed-effects models (CLMMs) with a logit link and flexible threshold parameters. Each model included the system as a fixed effect, with random intercepts and system-specific random slopes for both participants and questions. This specification accounts for inter-rater calibration differences and repeated-measure dependencies. Analyses were conducted at three levels: (i) Overall model (all metrics, all questions), (ii) Metric specific models, (iii) question-type specific models (query vs reasoning). Significance of interaction terms was evaluated via likelihood ratio tests and retained when statistically significant ($p < 0.05$).

Effect estimates are reported as odds ratios (ORs) quantifying the likelihood that React-to-Me received a higher rating relative to ChatGPT, with Wald-type 95% confidence intervals. One-sided p-values were derived from two-sided Wald tests by halving p-values when effect estimates favored React-to-Me (OR > 1).

**Nonparametric Validation.** To ensure robustness, model-based inferences were validated using distribution-free methods. Paired Wilcoxon signed-rank tests computed one-sided p-values and Hodges-Lehmann estimators with 95% confidence intervals for median score differences. Binomial sign tests were applied at two levels: (i) question-level comparisons across all evaluation pairs, and (ii) participant-level aggregated preferences.

**Effect Sizes.** Practical significance was quantified using: (i) Common-Language Effect Size (CLES), computed as strict win probability (excluding ties) and inclusive win probability (allocating half-credit to ties), with stratified bootstrap 95% confidence intervals (10,000 iterations, stratified by participant); (ii) paired Cliff's Delta; and (iii) Cohen's d.

**Multiple Testing Correction.** To control family-wise error rates, p-values were adjusted using the Holm step-down procedure within two families: (i) primary comparisons across three evaluation metrics, and (ii) secondary comparisons across two question types. Exploratory metric × question type stratifications were reported with uncorrected p-values.

**Power Analysis.** Post hoc achieved power was computed for each stratum using paired t-test approximations based on observed sample sizes and Cohen's d effect sizes ($\alpha = 0.05$, one-sided).

**Table S1 | Exploratory CLMM (Metric x Question Type)**

| Stratum | N (pairs) | OR [95% CI] | p (uncorr.) | CLES (strict) | Mean Δ (pts) | Wilcoxon p | Power |
|---|---|---|---|---|---|---|---|
| **Factual accuracy – Query** | 90 | 3.15 [1.52–6.53] | 0.0020 | 0.28 (0.19–0.37) | +0.27 | $1.16\times10^{-4}$ | 0.990 |
| **Factual accuracy – Reasoning** | 60 | 2.72 [0.94–7.91] | 0.065 | 0.27 (0.17–0.37) | +0.25 | 0.00287 | 0.900 |
| **Granularity – Query** | 90 | 2.97 [1.49–5.90] | 0.0019 | 0.41 (0.32–0.50) | +0.34 | $3.44\times10^{-4}$ | 0.975 |
| **Granularity – Reasoning** | 60 | 2.76 [0.85–9.01] | 0.092 | 0.45 (0.33–0.57) | +0.32 | 0.00608 | 0.832 |
| **Relational depth – Query** | 90 | 1.52 [0.74–3.14] | 0.26 | 0.29 (0.20–0.38) | +0.12 | 0.0810 | 0.405 |
| **Relational depth – Reasoning** | 60 | 2.57 [1.02–6.45] | 0.045 | 0.38 (0.28–0.50) | +0.30 | 0.00333 | 0.884 |

**Table S2 | Question-level sign tests**

| Stratum | N (pairs) | Wins | Losses | Ties | Win % | Mean Δ (pts) | Sign test p |
|---|---|---|---|---|---|---|---|
| **Overall** | 450 | 154 | 61 | 235 | 71.63 | 0.26 | $9.18\times10^{-11}$ ** |
| **Factual accuracy** | 150 | 41 | 9 | 100 | 82.00 | 0.26 | $2.81\times10^{-6}$ ** |
| **Level of granularity** | 150 | 64 | 26 | 60 | 71.11 | 0.33 | $3.83\times10^{-5}$ ** |
| **Relational depth** | 150 | 49 | 26 | 75 | 65.33 | 0.19 | $5.29\times10^{-3}$ ** |
| **Query** | 270 | 88 | 36 | 146 | 70.97 | 0.24 | $1.70\times10^{-6}$ ** |
| **Reasoning** | 180 | 66 | 25 | 89 | 72.53 | 0.29 | $1.01\times10^{-5}$ ** |
| **Factual accuracy – Query** | 90 | 25 | 5 | 60 | 83.33 | 0.27 | $1.62\times10^{-4}$ ** |
| **Factual accuracy – Reasoning** | 60 | 16 | 4 | 40 | 80.00 | 0.25 | $5.91\times10^{-3}$ ** |
| **Granularity – Query** | 90 | 37 | 14 | 39 | 72.55 | 0.34 | $8.85\times10^{-4}$ ** |
| **Granularity – Reasoning** | 60 | 27 | 12 | 21 | 69.23 | 0.32 | $1.19\times10^{-2}$ * |
| **Relational depth – Query** | 90 | 26 | 17 | 47 | 60.47 | 0.12 | $1.11\times10^{-1}$ |
| **Relational depth – Reasoning** | 60 | 23 | 9 | 28 | 71.88 | 0.30 | $1.00\times10^{-2}$ * |

**Table S3 | Participant-level sign tests**

| Stratum | n (participants) | Wins | Losses | Ties | Win % | Sign test p |
|---|---|---|---|---|---|---|
| **Overall** | 10 | 10 | 0 | 0 | 1.00 | $9.77\times10^{-4}$ ** |
| **Factual accuracy** | 10 | 9 | 1 | 0 | 0.90 | 0.0107 * |
| **Level of granularity** | 10 | 8 | 2 | 0 | 0.80 | 0.0547 |
| **Relational depth** | 10 | 9 | 1 | 0 | 0.90 | 0.0107 * |
| **Query** | 10 | 9 | 1 | 0 | 0.90 | 0.0107 * |
| **Reasoning** | 10 | 9 | 0 | 1 | 1.00 | 0.0020 ** |
| **Factual accuracy – Query** | 10 | 8 | 1 | 1 | 0.89 | 0.0195 * |
| **Factual accuracy – Reasoning** | 10 | 8 | 0 | 2 | 1.00 | 0.0039 ** |
| **Granularity – Query** | 10 | 7 | 2 | 1 | 0.78 | 0.0898 |
| **Granularity – Reasoning** | 10 | 8 | 1 | 1 | 0.89 | 0.0195 * |
| **Relational depth – Query** | 10 | 6 | 1 | 3 | 0.86 | 0.0625 |
| **Relational depth – Reasoning** | 10 | 8 | 1 | 1 | 0.89 | 0.0195 * |

#### 5 General User Satisfaction Survey

To complement the expert review, we conducted a general user survey to assess the real-world usability, interpretability, and scientific reliability of React-to-Me. The study protocol was approved by the University of Toronto Research Ethics Board (protocol #47192) and designed to collect structured, anonymous feedback from users interacting with the publicly deployed system at [https://reactome.org/chat](https://reactome.org/chat).

##### 5.1 Participant Recruitment and Eligibility

The survey aimed to capture a broad, international distributed sample of users across research sectors, and levels of biological expertise. Recruitment was conducted through local institutional bulletins and global online outreach via trusted scientific communities and networks. To encourage engagement from users with an active chatbot experience, a contextual prompt was embedded within the chat interface and triggered during typical usage. The survey was also accessible via the React-to-Me homepage for broader discovery. The goal was to ensure a diverse and representative participant pool reflective of the platform's intended user base.

Eligibility criteria required participants to be ≥18 years old, possess at least a bachelor's degree in a biology-related field, and have sufficient English proficiency to evaluate scientific content. Individuals with conflicts of interest (e.g., Reactome staff, close personal associates or active LLM developers) were excluded. Participation was voluntary, remote and uncompensated. As of September 2025, 27 participants had completed the full survey, with data collection ongoing since its launch on February 03, 2025.

##### 5.2 Survey Access and Procedure

Participants first reviewed an online informed consent form outlining the study's purpose, procedures, risks, and data handling. Those providing consent were invited to freely interact with the chatbot for approximately 15 minutes, followed by a 15–20 minute structured questionnaire.

The survey was designed to capture first hand impressions from users interacting with the live system, thereby reflecting authentic usage scenarios.

##### 5.3 Survey Design and Evaluation Metrics

The survey evaluated user experience across core performance metrics focusing on interpretability, usability, and perceived scientific quality.

The first component of the survey, Answer Quality and Usability, comprised ten Likert-scale questions evaluating key aspects of chatbot output and interaction. These included:

- Factual accuracy: alignment of chatbot responses with established biomedical knowledge.
- Relevance: how well responses addressed the user's intent and context .

- Clarity and conciseness: linguistic fluency and information density.
- Hyperlink and citation correctness: presence and validity of supporting references
- Logical consistency: coherence within and across conversational turns.
- Multilingual fluency: chatbot performance in languages other than English.
- Ease of use
- Overall satisfaction

These items were informed by existing literature on Human-AI and human-computer interaction studies, and refined through internal pilot testing

The Open-Ended Feedback section included three optional free-text prompts. Participants were encouraged to report any hallucinated or fabricated content, suggest specific areas for improvement, and describe concerns related to usability, accessibility, or scientific fidelity. These responses offered qualitative insights into edge cases and emergent usage patterns not captured by structured questions.

The last section, Demographics and Usage Context, collected optional non-identifying background information. Participants could indicate their geographic region, institutional type, field of research, education level in biology, preferred scientific languages, and Reactome usage frequency.

The full survey instrument, including item phrasing and response scales, is provided in Supplemental Appendix 2.

##### 5.4 Data Handling and Analysis

Survey responses were collected through Google Forms, exported locally, encrypted, and securely stored on institutional servers. Source data were deleted from the Google platform after export. No IP addresses, session metadata, or personally identifiable information (PII) were collected. Participants could withdraw at any point prior to data aggregation.

Quantitative responses were analyzed using descriptive statistics (means, standard deviations, and score distributions). Relationships among performance metrics were examined using pearson correlation coefficients to identify predictors of overall satisfaction.

To investigate whether domain familiarity influenced system evaluation, responses were further stratified by (i) self-reported education levels in biology (BSc in biology / interdisciplinary, Graduate level in biology, Beyond graduate level in biology) and (ii) Reactome familiarity and usage experience ( $< 1$ year vs. $\geq 1$ year).

Given the ordinal nature of responses and the modest sample size ($n = 27$), we employed two complementary nonparametric analysis approaches.

Ordinal analyses: responses were encoded as ordered categories ordered numerically(0 = needs improvement; 3 = works well). Experience-level comparisons (2 groups) used Mann-Whitney U tests; education-level comparisons (3 groups) used Kruskal-Wallis tests. Significant omnibus tests ($p < 0.05$) were decomposed via pairwise Mann-Whitney U tests with Bonferroni correction ($\alpha = 0.0167$ for three comparisons). Rank-biserial correlation quantified effect sizes.

Binary analyses: responses were binarized to "Works Well" vs. "Other" to test differences in satisfaction rates. Chi-square tests of independence were used for group comparisons, with Fisher's exact test applied when minimum expected cell counts fell below 5. Cramér's V quantified association strength (0.1 = small, 0.3 = medium, 0.5 = large).

Qualitative analysis: open-text responses were analyzed using inductive thematic analysis to identify recurring concerns, feature requests and patterns in user experience.

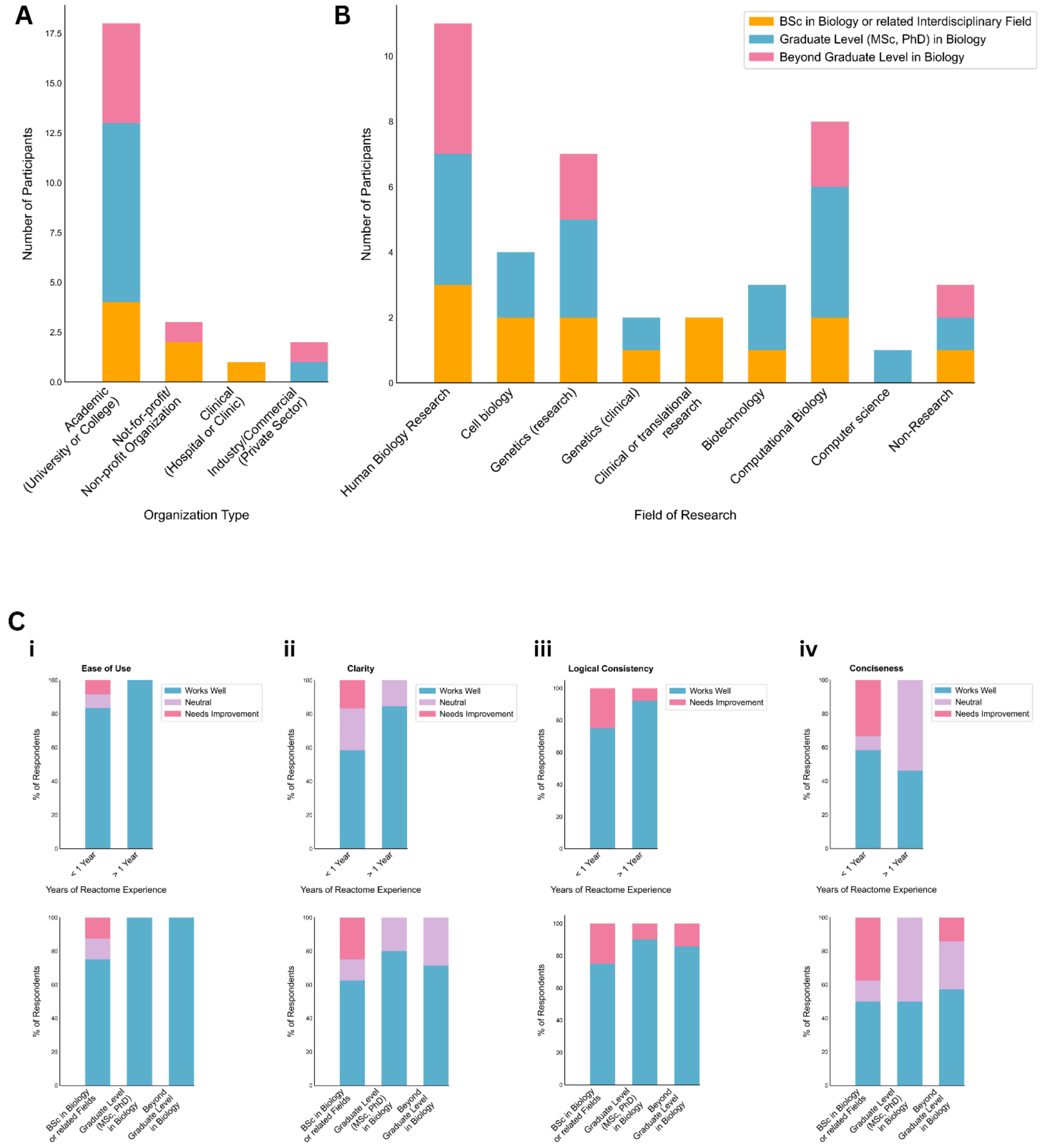


**Supplementary Figure 1 |**

**A) Respondent distribution by organization type (n=27).** Stacked bars show the education level composition within each category.

**B) Distribution by research field (n=27).** Human biology research most represented (41%), followed by computational biology (30%), cell biology (15%), and genetics (7%).

**C) Satisfaction ratings stratified by Reactome experience and education level across (i) ease of use, (ii) clarity, (iii) logical consistency, and (iv) conciseness**. Top row: comparison between users with <1 year versus ≥1 year Reactome experience. Bottom row: comparison across education levels (BSc/interdisciplinary, graduate level, beyond graduate level). Each panel shows percentage distributions across rating categories (Works Well/Neutral/Needs Improvement).
