## Supplemental Appendix 1 for "React-to-Me: A Conversational Interface for Interactive Exploration of the Reactome Pathway Knowledgebase"

### *Expert Evaluation of Chatbot Responses*

### *Survey Toolkit*

### Introduction:

***Welcome, and thank you for contributing your expertise.*** As a researcher in the biological sciences, your perspective is essential to understanding how effectively conversational AI can address complex, domain-specific questions. This study is part of an ongoing effort to explore the role of AI in supporting biomedical research and improving access to scientific knowledge.

Your insights will help evaluate the quality of AI-generated responses to biological questions, guide the development of more accurate and transparent tools, and contribute to the future of scientific information access in the life sciences.

### Study Objectives:

Structured biological databases like Reactome provide expert-curated knowledge of biological pathways, molecular mechanisms, and disease processes. However, these platforms can be difficult to navigate without significant technical expertise.

In contrast, large language models (LLMs) have made it easier to query complex subjects using natural language, often at the cost of domain specificity, accuracy, and transparency.

To bridge this gap, we've developed React-to-Me (https://reactome.org/chat), a conversational assistant grounded in Reactome content. This study evaluates whether React-to-Me provides more accurate, detailed, and mechanistically informed responses to biological questions than a general-purpose chatbot.

Your evaluations will play a key role in advancing AI tools for biomedical research, informing model refinement, benchmarking standards, and improving how scientists access and interact with complex biological information.

### Survey Structure:

The evaluation will be conducted using **Google Forms** and is designed to be intuitive, flexible, and aligned with your domain expertise. The full survey takes **approximately 2 hours to complete**, and you may pause and return at any time.

You will review **15 pairs of chatbot-generated responses** (30 responses in total). Each pair consists of two unlabeled answers to the **same question**, presented in random order to ensure unbiased evaluation.

- **Questions 1–5:** General Biology questions
  - Pre-selected and identical for all evaluators

- **Questions 6–15:** Field-specific questions.
  - You will select 10 questions from a dropdown menu.
  - Please choose questions that best match your expertise or research interests.

Your task is to evaluate each response **independently**, based on the criteria provided in the next section. Responses should be judged on their **intrinsic quality**, not by comparison to one another.

### Survey Instructions:

Before beginning the evaluation, please follow the steps outlined below:

#### Step 1: Review Evaluation Criteria

Please take a few minutes to review the evaluation criteria included in this package. This ensures your assessments are consistent with the standardized scoring system used across all evaluators.

- **Table 1 (p.3):** Defines the three evaluation dimensions you will use to assess each response
    - **Factual Accuracy,**
    - **Level of Granularity**
    - **Relational Depth** .
- **Table 2 (p.5-6):** Provides the 4-point grading rubric for each metric, with descriptive examples.

Please ensure you are familiar with both tables before proceeding. These criteria serve as the benchmark for scoring all responses.

#### Step 2: Access the Survey

Once you have reviewed the evaluation materials, open the survey using the link below:
[Google Form Link]

- The survey is hosted in Google Forms.
- You may complete it in multiple sittings; progress is automatically saved.
- All responses are anonymous and voluntary.

#### Step 3: Evaluate Responses:

1. **Understand the question:**
    - Take a moment to carefully review the biological topic being addressed.
2. **(Optional) Consult the background excerpt::**
    - For each question, a short excerpt from Voet & Voet, Biochemistry is provided for background.
    - This is intended to **refresh your memory,** not to serve as a gold standard or grading key.
    - Do **not** score responses against this reference. Use it only for **optional context**.
3. **Evaluate each response independently:**
    - Do not compare responses to one another.
    - Evaluate each one **on its own merits** using the criteria from **Table 1 and Table 2**.

### Contact Information:

If you have any questions or need assistance while completing the survey, please feel free to contact me at!

**Table 1: Description of three dimensions: factual accuracy, level of granularity, and relational depth**

| Metric | Description |
|---|---|
| **Factual Accuracy** | ● All claims are factually correct, mechanistically sound, and supported by current **scientific knowledge** or the reference answer.<br>● Interpretable and trustworthy by **both** expert and non-expert readers. |
| **Level of Granularity** | ● Demonstrates expert-level precision in selecting and integrating relevant biological entities.<br>● Each mentioned gene, protein, pathway, reaction, or disease is **directly relevant** and **necessary** to fully answer the question. |
| **Relational Depth** | ● Provides nuanced, **mechanistic insight** into how identified elements function and relate to each other within a biological system.<br>● Clearly explains **interactions** (e.g., causal pathways, regulatory effects, signaling dynamics) and **contextual relevance**.<br>● Demonstrates systems thinking. |

**Here is a simple example, to understand the general concept:**

**Question: "Why do we need sleep?"**

**1. High Factual Accuracy, Low Level of Granularity, Low Relational Depth:**

- Sleep helps the body rest and recover. Without it, we feel tired and can't function properly.

***Commentary:*** *Factually accurate, but doesn't mention specific systems or processes. Overly broad, vague and aimed at a non-expert audience.*

**2. High Accuracy,  High Level of Granularity, Low Relational Depth:**

- Sleep is regulated by circadian rhythms and involves stages like REM and non-REM sleep.
- During sleep, the body releases melatonin, restores neurotransmitter levels, and activates processes like memory consolidation.

***Commentary:*** *Includes specific, relevant terms (circadian rhythms, REM, melatonin), and correct facts. However, it presents them as a loose collection rather than a coherent, mechanistic explanation. Granular, but not deeply connected.*

**3. High Accuracy, Low Level of Granularity, High Relational Depth:**

- Sleep is a coordinated biological state that allows the brain and body to perform essential maintenance functions.
- It supports neural stability, regulates metabolism, and maintains immune balance.
- These processes work together to preserve long-term physiological health.

***Commentary:*** *Provides a strong systems-level explanation — why sleep matters, what it's doing, how it connects across systems — but without naming specific hormones, brain regions, or pathways. High depth, low granularity.*

**4. Low Accuracy, High Level of Granularity, High Relational Depth:**

- During sleep, neurons regenerate and multiply to replace those lost during wakefulness.
- The pineal gland signals the brain to enter REM cycles through cortisol release, which activates glymphatic flow and triggers memory storage in the cerebellum.
- Over time, insufficient sleep causes neurons to stop firing, leading to permanent brain shutdown.

***Commentary:*** *Response is detailed and presents a seemingly coherent mechanistic narrative, but it is factually inaccurate on multiple fronts. The language and structure mimic expert discourse, yet key claims are biologically false or misleading.*

**5. High Accuracy, High Level of Granularity, High Relational Depth:**

- Sleep is regulated by circadian signals from the suprachiasmatic nucleus and melatonin release.
- During sleep, the brain clears waste via the glymphatic system, consolidates memory through hippocampal-neocortical activity, and restores neurotransmitter balance.
- At the same time, immune function is recalibrated and metabolic pathways are reset.
- These coordinated processes maintain cognitive performance, physiological stability, and long-term health

**Table 2: Grade scale, with examples of different levels:**
**Example Question: *"Explain the role of the PI3K/AKT pathway in cancer development and progression."***

| Grade Scale | Level 1: Poor (0-25%) | Level 2: Fair (26-50%) | Level 3: Good (51-75%) | Level 4: Excellent (76-100%) |
|---|---|---|---|---|
| **Factual Accuracy** | | | | |
| **Description** | Contains multiple major factual errors. Claims are incorrect, contradict established scientific understanding, or miscommunicate underlying biology.<br><br>Would mislead both experts and non-experts. | Includes some accurate statements, but central elements are incorrect or presented in ways that distort underlying biology.<br><br>Misleading to non-experts and problematic for expert interpretation. | Largely factually correct. May contain minor inaccuracies or subtle mechanistic errors.<br><br>Would generally be recognized as valid by experts, though some phrasing could mislead non-experts. | All claims are factually correct, mechanistically sound, and supported by current **scientific knowledge** or the reference answer.<br><br>Interpretable and trustworthy by both expert and non-expert readers. |
| **Example Answer** | PI3K/AKT repairs cancer by fixing DNA. | PI3K promotes tumour growth by turning on mitosis genes directly. | PI3K causes mitosis directly, leading to cancer. | Loss of PTEN activates PI3K/AKT signaling, promoting tumor growth |
| **Pitfall** | *Scientifically Incorrect* | *Misrepresents causal relationships; superficially plausible but conceptually flawed.* | *Oversimplified mechanism that **overstates** direct causality. Mechanistically misleading* | |
| **Level of Granularity** | | | | |
| **Description** | Response lacks biological specificity. No clearly identified biological entities (e.g., genes, proteins, pathways) are present.<br><br>Uses vague language or generalizations that could apply to any biological process. | Identifies a few relevant biological entities though may miss important details or include irrelevant content.<br><br>Terms may be technically correct but insufficient to meaningfully answer the question | Identifies multiple contextually relevant biological entities. Not all included details are essential or well-targeted.<br><br>Demonstrates domain fluency but **lacks expert-level selectivity**. | Each mentioned gene, protein, pathway, reaction, or disease is **directly relevant** and **necessary** to fully answer the question.<br><br>Demonstrates expert-level precision in selecting and integrating relevant biological entities. |
| **Example Answer** | Something in the cell goes wrong and cancer starts. | AKT activation promotes cell survival and division, which can support cancer progression. | Activation of PI3K leads to downstream effects on AKT, mTORC1, and FOXO transcription factors, regulating processes such as cell survival, metabolism, growth, and autophagy, which are often altered in cancer. | Loss of PTEN increases PIP3 levels, activating AKT through membrane recruitment. Activated AKT promotes cell survival and proliferation, contributing to tumorigenesis. |
| **Pitfall** | *Lacks specificity; Uses no relevant terminology or clearly defined concepts.* | *Identifies **one** correct, relevant protein; introduces some broadly relevant domain-specific terms.* | *The granularity is excessive, leading to reduced focus.* | *Focused, biologically optimal selection of entities that directly supports the question.* |

| Relational Depth | | | | |
|---|---|---|---|---|
| **Description** | Response is vague or superficial. Offers no meaningful biological explanation of relationships between components. No indication of underlying mechanisms or contextual understanding. | Provides a general explanation of key concepts. May mention functions (e.g., proliferation, survival) but lacks specificity about how elements influence each other. Relationships may be correct but shallow or one-dimensional. | Demonstrates mechanistic understanding of how entities function in relation to each other. Describes directional or regulatory effects with some clarity. May omit broader system interactions (e.g., cross-talk, feedback, temporal dynamics). | Provides nuanced, **mechanistic insight** into how identified elements function and relate to each other within a biological system. Clearly explains interactions (e.g., causal pathways, regulatory effects, signaling dynamics) and contextual relevance. Demonstrates systems thinking. |
| **Example Answer** | It helps cells grow, so cancer happens. | AKT helps cells survive, and if it's always active, that can lead to cancer. | Loss of PTEN leads to unchecked PI3K/AKT activity, promoting cell survival and growth, which contributes to tumor progression. | Loss of PTEN leads to PIP3 and constitutive recruitment and activation of AKT at the plasma membrane. Activated AKT promotes cell survival and proliferation while modulating metabolic pathways and suppressing immune responses. These changes collectively contribute to a tumor-permissive microenvironment. |
| **Pitfall** | *Oversimplified and vague; no biological terms or mechanisms.* | *Identifies the pathway and* ***surface-level*** *function; lacks causality, mechanistic detail or systems-level interaction* | *Correct and causal, but limited to linear or isolated pathways; does not capture system-level complexity or contextual dependencies.* | ***Mechanistic*** *detail, protein function, signaling flow, and disease context all clearly explained.* |
