## Supplemental Appendix 2 for "React-to-Me: A Conversational Interface for Interactive Exploration of the Reactome Pathway Knowledgebase"

### React-To-Me Chatbot Response Quality General Survey Informed Consent Overview

#### Purpose of study:

This study is conducted by researchers at the Ontario Institute for Cancer Research.

The purpose of this study is to evaluate the effectiveness of a new AI-driven conversational interface designed to improve accessibility and usability of the Reactome database of biological pathways. This study aims to evaluate if our AI chatbot can enhance access to the Reactome database by providing accurate and contextually relevant information with fewer AI-generated errors. Through this study, we aim to gather insights into the React-To-Me chatbot's usability, its ability to enhance user understanding of complex biological pathways, and its overall potential to improve the research experience for a diverse range of users. You must be at least 18 years of age to participate.

#### Procedures:

You will have the opportunity to interact with the chatbot, and we encourage you to **use your preferred or native language**, as the chatbot supports multiple languages. After the interaction, you will be asked to complete a questionnaire about your experience with the React-To-Me AI Chatbot Prototype. This should take approximately 15-20 minutes. The questions will focus on your general interaction with the chatbot, with optional sections regarding your educational background and familiarity with biological pathways or the Reactome database. At the end of the questionnaire you will be asked if you would like to meet with the researchers for a follow-up interview which will take place by videoconferencing.

#### Potential Risks and Benefits:

It is unlikely that you will experience any risks or discomforts beyond what you would experience when interacting with any other AI chatbot. It is possible, but unlikely, that the AI chatbot will make statements that are inflammatory or inappropriate. In the event that this does occur, your participation will help us to identify interactions that trigger this behavior and improve the AI chatbot to avoid such language in the future.

We do not expect you to benefit from the study, though it is possible that the interactions improve your understanding of the Reactome knowledgebase contents, or general molecular biology concepts. Your participation helps us evaluate and improve a new AI chatbot that benefits the scientific community.

#### Compensation:

There is **no compensation** for completing the general questionnaire.

#### Confidentiality:

The data collected is anonymous, and the data you provide will be anonymized and aggregated. No personally identifiable data is collected. Any public report will not include any information that will make it possible to identify you. Any quotations or testimonials included in a public report, with your permission, will be anonymized.

#### Voluntary Participation:

Your participation in the study is **voluntary**. You may choose to end your participation at any time without penalty. If you choose to withdraw from the study, we will remove your responses to the questionnaire from the study database. However, if your responses were included in published aggregate statistics, then we will not be able to eliminate your contribution to the aggregate data. To request withdrawal from the study, send an email to:

#### Questions or Concerns:

Please take your time reading this form and contact the researchers to ask questions if there is anything you do not understand. If you have questions or concerns, you may contact the Reactome Outreach Coordinator Dr. Nancy Li at
React-To-Me chatbot

### React-To-Me Chatbot Response Quality General Survey Questions

#### Consent Question:

1. I have carefully read and fully understood the details provided in the consent form, and I willingly agree to participate in this study.
    a. Yes
    b. No

#### General Questions:

**Purpose:**
This section is designed to gather your feedback on the performance of the React-To-Me chatbot. Please reflect on your **recent interaction** with the chatbot when providing your responses. Your feedback will help us understand how well the chatbot meets your expectations.

**What to Expect:**
You will be asked to **rate various aspects of your experience**, including how accurate and relevant the chatbot's answers were, how concise and clear the responses felt, and how easy the chatbot was to use. Additionally, questions about the chatbot's consistency, citation accuracy, and support for languages other than English will help us ensure its usability across different contexts.

<u>Answer Quality</u>

1. **Accuracy:**How accurate were the answers provided by the chatbot?
    a. Very Accurate
    b. Accurate
    c. Neutral
    d. Inaccurate
    e. Very Inaccurate
2. **Conciseness:** How concise were the answers?
    a. Overly brief
    b. Appropriately concise
    c. Somewhat Lengthy
    d. Overly Lengthy
3. **Relevance:** How relevant were the answers to your questions?
    a. Highly Relevant
    b. Relevant
    c. Moderately Relevant
    d. Slightly Relevant

    e. Not Relevant

4. **Clarity:** Were the answers clear and easy to understand?
    a. Always
    b. Sometimes
    c. Never

5. **Citation:** Were the provided links correct?
    a. Always
    b. Sometimes
    c. Never

6. **Consistency:** Did the chatbot provide logically consistent answers throughout your interactions?
    a. Yes
    b. No

7. If your native / preferred language(s) for science is not English, did you attempt to converse with the chatbot in your native language?
    a. Yes
    b. No

8. If you interacted with the chatbot in a language other than English, how would you rate its fluency in that language? Does the chatbot support your preferred language(s)?
    a. Very fluent
    b. Moderately fluent
    c. Somewhat fluent
    d. Not very fluent
    e. Not fluent at all

<u>User Satisfaction:</u>

9. **Ease of Use:** How easy was it to use the chatbot?
    a. Very Easy
    b. Easy
    c. Neutral
    d. Difficult
    e. Very Difficult

10. **Overall Satisfaction:** How satisfied are you with your overall experience using the chatbot?
    a. Very Satisfied
    b. Satisfied
    c. Neutral
    d. Dissatisfied
    e. Very Dissatisfied

11. How frequently do you use the Reactome Knowledgebase to learn about biological processes?
    a. Daily
    b. Weekly
    c. Monthly

d. Rarely

12. How frequently do you use the UniProt Knowledgebase to learn about biological processes?
    a. Daily
    b. Weekly
    c. Monthly
    d. Rarely

13. How frequently do you use the Alliance of Genome Resources Knowledgebase to learn about biological processes?
    a. Daily
    b. Weekly
    c. Monthly
    d. Rarely

14. How frequently do you use the UniProt Knowledgebase to learn about biological processes?
    a. Daily
    b. Weekly
    c. Monthly
    d. Rarely

15. **Comparison:** How does the chatbot compare to other sources of information you have consulted to learn about biological processes (e.g., UniprotKB, Reactome KB, Alliance, Pubmed, Wikipedia, Google)?
    a. Much Better
    b. Better
    c. About the same
    d. Worse
    e. Much worse

User Engagement:

1. How satisfied are you with the chatbot's ability to maintain an engaging conversation?
    a. Very Satisfied
    b. Satisfied
    c. Moderately Satisfied
    d. Slightly Satisfied
    e. Not Satisfied

2. Based on your experience today, Do you see yourself using this chatbot regularly?
    a. Yes
    b. Maybe
    c. No

Open-ended questions:

1. Do you have any additional comments or suggestions for improving the chatbot?

2. If you have any concerns or encountered any issues, please describe them here.

#### 2.4 Demographic Questions:

The following are standard questions that allow researchers to determine how representative the group of participants in a study is of the general population. These questions are voluntary.

1. How long have you been using Reactome resources?
    a. Never before today
    b. Less than 6 months
    c. 6 months to 1 year
    d. 1 year to five years
    e. More than 5 years

2. Which of these statements best describe your usage of the Reactome website or tools?
    a. I use Reactome on a daily basis
    b. I use Reactome on a weekly basis
    c. I use Reactome on a monthly basis
    d. I use Reactome rarely

3. Which of these statements best describe your highest education level?
    a. I received a formal education at or beyond graduate-level in biology (ex. post-doc, staff scientist, PI)
    b. I received a formal education at graduate-level in biology
    c. I received a formal education at BSc-level in biology
    d. I received some formal training in a biology or interdisciplinary field
    e. Other (please specify)

4. What part of the world do you work in?
    - AF - Afghanistan
    - AL - Albania
    - DZ - Algeria
    - AD - Andorra
    - AO - Angola
    - AG - Antigua and Barbuda
    - AR - Argentina
    - AM - Armenia
    - AU - Australia
    - AT - Austria
    - AZ - Azerbaijan
    - BS - Bahamas
    - BH - Bahrain
    - BD - Bangladesh
    - BB - Barbados
    - BY - Belarus
    - BE - Belgium
    - BZ - Belize
    - BJ - Benin
    - BT - Bhutan
    - BO - Bolivia
    - BA - Bosnia and Herzegovina
    - BW - Botswana
    - BR - Brazil
    - BN - Brunei Darussalam

- BG - Bulgaria
- BF - Burkina Faso
- BI - Burundi
- CV - Cabo Verde
- KH - Cambodia
- CM - Cameroon
- CA - Canada
- CF - Central African Republic
- TD - Chad
- CL - Chile
- CN - China
- CO - Colombia
- KM - Comoros
- CG - Congo
- CR - Costa Rica
- CI - Côte D'Ivoire
- HR - Croatia
- CU - Cuba
- CY - Cyprus
- CZ - Czechia
- CD - Democratic Republic of the Congo
- DK - Denmark
- DJ - Djibouti
- DM - Dominica
- DO - Dominican Republic
- EC - Ecuador
- EG - Egypt
- SV - El Salvador
- GQ - Equatorial Guinea
- ER - Eritrea
- EE - Estonia
- SZ - Eswatini
- ET - Ethiopia
- FJ - Fiji
- FI - Finland
- FR - France
- GA - Gabon
- GM - Gambia
- GE - Georgia
- DE - Germany
- GH - Ghana
- GR - Greece
- GD - Grenada
- GT - Guatemala
- GN - Guinea
- GW - Guinea Bissau
- GY - Guyana
- HT - Haiti
- HN - Honduras
- HU - Hungary
- IS - Iceland

- IN - India
- ID - Indonesia
- IR - Iran
- IQ - Iraq
- IE - Ireland
- IL - Israel
- IT - Italy
- JM - Jamaica
- JP - Japan
- JO - Jordan
- KZ - Kazakhstan
- KE - Kenya
- KI - Kiribati
- KW - Kuwait
- KG - Kyrgyzstan
- LA - Laos
- LV - Latvia
- LB - Lebanon
- LS - Lesotho
- LR - Liberia
- LY - Libya
- LI - Liechtenstein
- LT - Lithuania
- LU - Luxembourg
- MG - Madagascar
- MW - Malawi
- MY - Malaysia
- MV - Maldives
- ML - Mali
- MT - Malta
- MH - Marshall Islands
- MR - Mauritania
- MU - Mauritius
- MX - Mexico
- FM - Micronesia
- MC - Monaco
- MN - Mongolia
- ME - Montenegro
- MA - Morocco
- MZ - Mozambique
- MM - Myanmar
- NA - Namibia
- NR - Nauru
- NP - Nepal
- NL - Netherlands
- NZ - New Zealand
- NI - Nicaragua
- NE - Niger
- NG - Nigeria
- KP - North Korea
- MK - North Macedonia

- NO - Norway
- OM - Oman
- PK - Pakistan
- PW - Palau
- PA - Panama
- PG - Papua New Guinea
- PY - Paraguay
- PE - Peru
- PH - Philippines
- PL - Poland
- PT - Portugal
- QA - Qatar
- MD - Republic of Moldova
- RO - Romania
- RU - Russian Federation
- RW - Rwanda
- KN - Saint Kitts and Nevis
- LC - Saint Lucia
- VC - Saint Vincent and the Grenadines
- WS - Samoa
- SM - San Marino
- ST - Sao Tome and Principe
- SA - Saudi Arabia
- SN - Senegal
- RS - Serbia
- SC - Seychelles
- SL - Sierra Leone
- SG - Singapore
- SK - Slovakia
- SI - Slovenia
- SB - Solomon Islands
- SO - Somalia
- ZA - South Africa
- KR - South Korea
- SS - South Sudan
- ES - Spain
- LK - Sri Lanka
- SD - Sudan
- SR - Suriname
- SE - Sweden
- CH - Switzerland
- SY - Syrian Arab Republic
- TJ - Tajikistan
- TZ - Tanzania
- TH - Thailand
- TL - Timor-Leste
- TG - Togo
- TO - Tonga
- TT - Trinidad and Tobago
- TN - Tunisia
- TR - Turkey

- TM - Turkmenistan
- TV - Tuvalu
- UG - Uganda
- UA - Ukraine
- AE - United Arab Emirates
- GB - United Kingdom
- US - United States of America
- UY - Uruguay
- UZ - Uzbekistan
- VU - Vanuatu
- VE - Venezuela
- VN - Viet Nam
- YE - Yemen
- ZM - Zambia
- ZW - Zimbabwe
- Other (please specify)

5. What type of organization do you work for or study at?
    a. Academic (University or College)
    b. Not-for-profit/Non-profit Organization
    c. Hospital-based Research Centre/Institute
    d. Clinical (Hospital or Clinic)
    e. Industry/Commercial (Private Sector)
    f. Government (Public Sector)
    g. Independent Researcher
    h. Other (Please specify)

6. What term(s) best describe the type of research you perform?
    a. I am not a researcher
    b. Basic research (human)
    c. Basic research (model organism)
    d. Biotechnology
    e. Bioinformatics or computational biology
    f. Chemistry or chemoinformatics
    g. Clinical or translational research
    h. Computer science
    i. Genetics (research)
    j. Genetics (clinical)
    k. Pharmacology
    l. Cell biology
    m. Systems biology
    n. Other (Please specify)

7. What is your preferred language(s) for science?
    a. English
    b. French
    c. Spanish
    d. Portuguese
    e. Chinese
    f. Korean
    g. Japanese

h. Hindi
i. Urdu
j. Farsi
k. Arabic
l. German
m. Russian
n. Italian
o. Dutch
p. Turkish
q. Thai
r. Vietnamese
s. Polish
t. Ukrainian
u. Greek
v. Hebrew
w. Bengali
x. Tamil
y. Malay/Indonesian
z. Swahili
aa. Other(please specify)

2.5 Future Participation:

1. Would you be willing to be contacted in the future to participate in follow-up interviews or testing sessions to further refine the LLM?

    a. Yes
    b. No
